# Suppression of PP2A-B56α Drives EMT in EGFR Mutant Non-Small Cell Lung Cancer

**DOI:** 10.1101/2025.07.14.664751

**Authors:** Brittany N. Heil, Garima Baral, Claire M. Pfeffer, Mei B. Bahler, Anna K. Darling, Sydney J. Clifford, Whitney Smith-Kinnaman, Kasi Hansen, Emma H. Doud, Gaganpreet K. Mall, Aaron N. Hata, Nicole L. Anderson, Matthew R. Olson, Brittany L. Allen-Petersen

## Abstract

Lung cancer is the leading cause of cancer-related deaths in the United States and ∼50% of these patients present with metastatic disease at diagnosis. Epithelial-to-Mesenchymal Transition (EMT) is an important initiating step in the metastatic cascade that allows cells to acquire the migratory and invasive phenotypes necessary for dissemination. The transcriptional reprogramming that takes place during EMT has been well described in multiple cancer types; however, the posttranslational regulatory mechanisms that govern EMT are poorly understood. Protein Phosphatase 2A (PP2A) is serine/threonine (ser/thr) phosphatase that accounts for 50% of cellular ser/thr phosphatase activity and is critically important in regulating signaling homeostasis. PP2A dysregulation has been implicated in cell state regulation, EMT, and metastasis, but the roles of individual PP2A complexes are poorly understood. Our data indicate that suppression of the specific PP2A complex, PP2A-B56α, results in decreased expression of epithelial markers and increased expression of mesenchymal markers consistent with EMT. These molecular changes are associated with migratory and invasive phenotypes both *in vitro* and *in vivo*. Furthermore, these migratory phenotypes can be rescued with B56α overexpression. Together, these findings implicate B56α as a key regulator of cellular plasticity and highlight the dynamic nature by which PP2A-B56α posttranslationally regulates NSCLC EMT.

## INTRODUCTION

Lung cancer is the leading cause of cancer-related deaths worldwide and has an overall 5-year survival rate of approximately 26%, which plummets to just 9% with metastatic disease [1]. In Non-Small Cell Lung Cancer (NSCLC), the most common type of lung cancer, about 50% of patients have metastatic disease at the time of diagnosis [2]. During tumor progression, cancer cells acquire metastatic capabilities by undergoing an epithelial-to-mesenchymal transition (EMT). This unique form of cellular plasticity results in aggressive tumor phenotypes and is associated with broad therapeutic resistance [3–5]. Given the impact of EMT on tumor progression, understanding the mechanisms that regulate this process is essential for improving patient outcomes.

EMT is a dynamic and reversible process that is associated with reduced cell-cell junctions, acquired front-back polarity, and increased migratory and invasive capabilities, which allow cells to survive dissemination and travel to distant metastatic sites. Aberrant activation or expression of EMT-inducing transcription factors (EMT-TFs) (e.g. ZEB1, TWIST, and SNAIL) potently drives EMT in a wide variety of solid tumors [6–9]. In addition to these transcriptional programs, it is now appreciated that numerous posttranslational mechanisms play key roles in the regulation of EMT, including phosphorylation. Cancer cells can undergo EMT in response to a wide variety of signals, including therapeutic stress, without the need for additional oncogenic mutations, suggesting that posttranslational signaling mechanisms play a key role in driving EMT [10,11]. Accordingly, several EMT-TFs are activated downstream of kinase signaling cascades and direct phosphorylation of these transcription factors can impact their activity by altering both stability and subcellular localization [12]. Similarly, the localization of proteins involved in cellular adhesion are directly impacted by phosphorylation, suggesting that transcriptional and posttranslational regulatory mechanisms work together to maintain epithelial cell fates. These studies highlight the impact of non-genomic signaling mechanisms on cellular plasticity; however, our understanding of the posttranslational events that govern EMT remain limited. Additionally, as the analysis of patient tumors rarely includes posttranslational phosphorylation events, the dysregulation of posttranslational mechanisms may be underestimated in metastatic patient populations.

Protein phosphatases function as key gatekeepers of cellular plasticity, providing essential regulation of intracellular signaling cascades. Protein Phosphatase 2A (PP2A) is a heterotrimeric serine-threonine phosphatase that has been identified as a vital tumor suppressor in many cancer types and negatively regulates many of the downstream effectors of the Epidermal Growth Factor Receptor (EGFR) pathway, the most common genetic alteration in NSCLC [13]. The PP2A holoenzyme is composed of 3 subunits: the scaffolding “A” subunit, the catalytic “C” subunit, and 16 regulatory “B” subunits that determine substrate specificity. PP2A has been implicated in the direct regulation of many cell-cell junction proteins including desmosomes, E-cadherin, and β-catenin [14]. Pharmacological inhibition or genetic loss of total PP2A activity disrupts these junctions and contributes to metastasis in several cancer types; however, the contribution of individual B subunits to the regulation of cancer cell signaling pathways is largely unknown. Using shRNA knockdown screens, Sablina et al. determined that the specific PP2A subunit, B56α, displays potent tumor suppressor capabilities [15]. Similarly, genetic loss of B56α results in low-penetrance tumor formation and increased stem-like properties after long latency [16]. In melanoma patients, expression of B56α is significantly reduced in metastatic versus primary tumors, implicating a role for B56α in tumor dissemination [17]. Recently, our lab demonstrated that loss of B56α in combination with oncogenic KRAS^G12D^ expression increased chromatin accessibility at EMT related genes and exacerbated cancer-associated cell fate transitions [18]. Together, these findings suggest that PP2A-B56α functions as a key regulator of cancer cell plasticity; however, the contribution of this B subunit to NSCLC EMT has not been assessed. Therefore, we hypothesize that PP2A-B56α inhibits NSCLC cancer cellular plasticity and suppresses the posttranslational activation of EMT signaling programs.

Here, we show that reduced B56α expression correlates with decreased overall NSCLC patient survival. In primary EGFR mutant NSCLC patient-derived cell lines, B56α expression is correlated with expression of the epithelial cell marker, E-cadherin. Similarly, suppression of B56α results in a morphological shift from an epithelial to a mesenchymal cell state consistent with EMT. This cell state change is associated with a loss of epithelial cell-cell junctions and a corresponding increase in mesenchymal marker expression. Beyond these markers, the B56α-dependent EMT phenotype is associated with large-scale proteomic and phosphoproteomic changes indicative of broad cellular rewiring and aberrant pathway activation. In line with the role of EMT in metastasis, B56α knockdown cells have a significant increase in migratory and invasive capacity both *in vitro* and *in vivo*. These phenotypes can be rescued with overexpression of B56α highlighting the dynamic and reversible nature of this cellular plasticity. Together our findings support a critical role for PP2A-B56α in the regulation of EMT cell state dynamics and metastasis in EGFR-driven NSCLC.

## RESULTS

### PP2A-B56α plays a critical role in NSCLC tumor progression

To identify the relationship between PP2A-B56α expression and NSCLC patient outcomes, we analyzed publicly available genomic data from NSCLC patients. We found that B56α (*PPP2R5A*) mRNA expression is significantly decreased in NSCLC tumors compared to adjacent normal tissue (Figure 1A). Similarly, low B56α (*PPP2R5A*) mRNA expression or high mRNA expression of Cancerous Inhibitor of PP2A (CIP2A, *KIAA1524*), an endogenous inhibitor of the B56 family of subunits, correlated with decreased overall patient survival (Figure 1B, 1C) [19]. We and others have previously shown that B56α directly dephosphorylates the oncoprotein, c-MYC (MYC), at Serine62 (S62) which drives MYC degradation [18,20–23]. Recently, we established that activation of the mitogen activated protein kinase (MAPK) pathway downstream of EGFR/RAS signaling leads to increased CIP2A expression, decreased PP2A-B56α activity, and increased phosphorylation of S62 MYC [18]. These signaling changes were associated with increased cellular plasticity and EMT gene programs. Based on these findings, we sought to determine the expression levels of B56α, E-cadherin (epithelial marker), and Vimentin (Mesenchymal marker) in primary NSCLC patient derived cell lines. These lines were generated from patients with NSCLCs harboring *EGFR* exon 19 deletions (activating alteration) either before or after treatment with EGFR tyrosine kinase inhibitors (Figure 1D)[24,25]. The lines exhibited diverse morphological characteristics ranging from epithelial to mesenchymal, and had varied expression of E-cadherin and Vimentin. Using quantitative RT-PCR (qRT-PCR) we determined that the expression B56α is positively correlated with E-cadherin (*p<0.05*)(Figure 1E) and trended with decreased Vimentin expression (*p=0.226*)(Figure 1F), suggesting that PP2A-B56α may function to sustain an epithelial cell identity in NSCLC.

**Figure 1:**
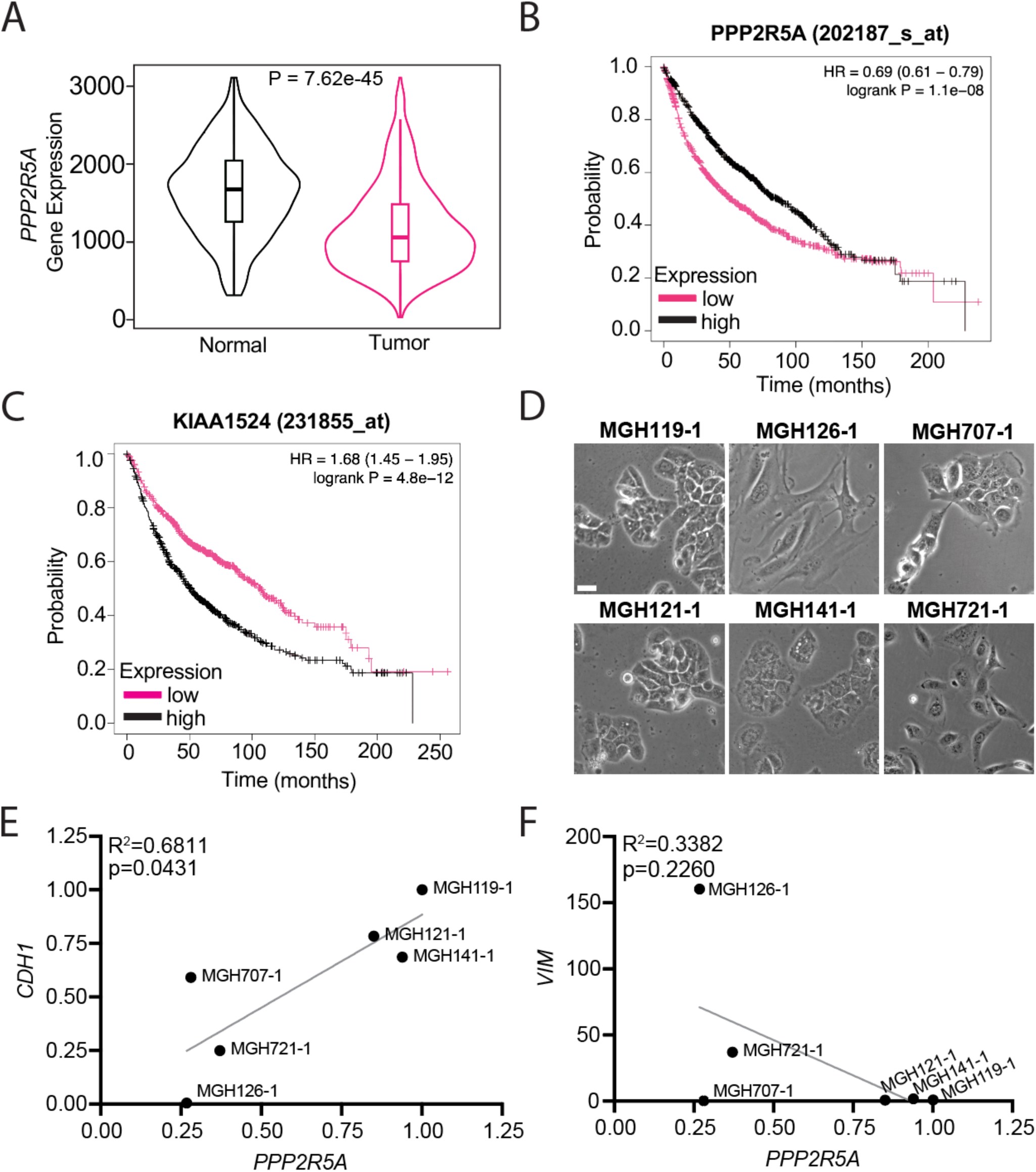
Low B56α expression correlates with poor prognosis in NSCLC. A) Normal and tumor tissue from lung cancer patient analyzed by RNA sequencing for B56α (*PPP2R5A*) expression, showing decreased expression in tumor tissue compared to normal adjacent tissue. B) Survival curve of lung cancer patients with low or high expression of B56α, with low expression of B56α correlating with decreased survival. C) Survival curve of lung cancer patients with low or high expression CIP2A (*KIAA1524*), with high expression of CIP2A correlating with decreased survival. KMPlotter was used to analyze panels A-C. D) Representative brightfield images of primary patient cell lines from NSCLC patients (scale bar = 100μm). E) Correlation of the mRNA expression of E-cadherin (*CDH1*) and B56α in a panel of NSCLC primary patient cell lines (n=3 biological replicates). F) Correlation of the mRNA expression of Vimentin (*VIM*) and B56α in a panel of NSCLC primary patient cell lines (n=3 biological replicates).

### PP2A-B56α suppression results in morphological and molecular changes associated with EMT

To determine the impact of PP2A-B56α suppression on NSCLC tumor phenotypes, we first knocked down the B56α subunit of PP2A using two independent shRNAs (shB56α1 and shB56α2) in human NSCLC cell lines with EGFR exon 19 deletion (EGFR del19) (Supplemental Figure 1A). Both H1650 and HCC827 shB56α cells exhibited decreased epithelial “cobblestone” morphology and increased front-back polarity compared to a scrambled shRNA control (shSCR) (Figure 2A, Supplemental Figure 1B). To determine if these morphological changes were associated with a loss in epithelial identity, expression of E-cadherin and Vimentin was analyzed in shB56α cells. B56α knockdown resulted in a significant decrease in E-cadherin mRNA and protein expression compared to shSCR controls (Figure 2B). H1650 and HCC827 shB56α cells were then fixed and DAPI, Vimentin, and E-cadherin were analyzed by immunofluorescent (IF) imaging. H1650 shB56α cells displayed a drastic loss of E-cadherin with a corresponding increase in Vimentin expression (Figure 2C). Similar results were found HCC827 shB56α cells albeit to a lesser degree, potentially indicative of a partial EMT phenotype (Figure 2D). Together, these results suggest that PP2A-B56α functions to maintain cellular epithelial identity, with inhibition of PP2A-B56α promoting a mesenchymal cell state.

**Figure 2:**
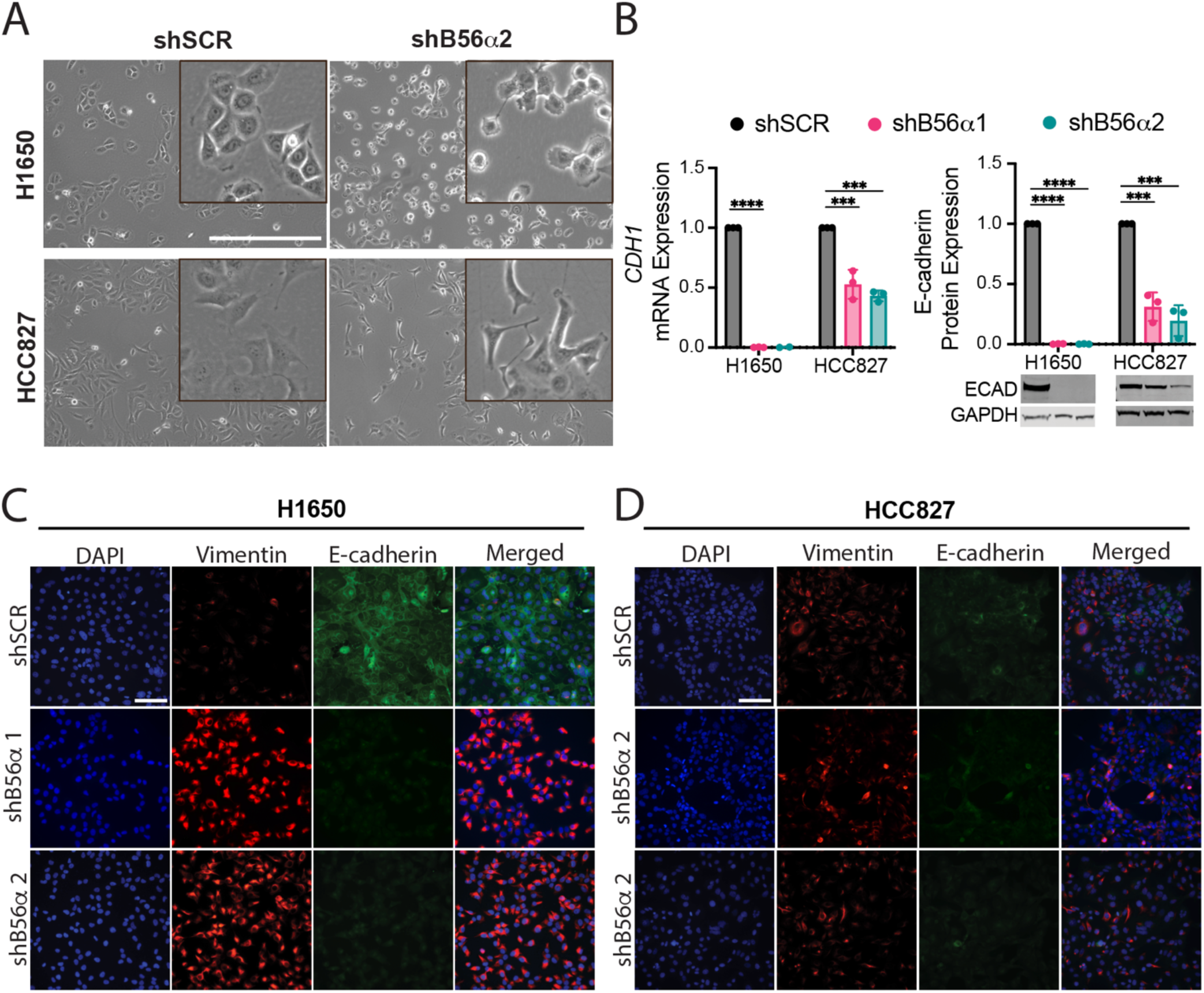
Knockdown of B56α drives EMT marker expression. A) Representative brightfield images of H1650 and HCC827 cell lines with shRNA mediated knockdown of B56α (KD) compared to shSCR control (scale bar=500μm). Inset denotes digital zoom. B) mRNA and protein expression of E-cadherin in H1650 and HCC827 KD cell lines compared to shSCR control (n=3 biological replicates) C) Representative immunofluorescent images from 3 biological replicates of H1650 KD and D) HCC827 KD cell lines with DAPI, Vimentin, and E-cadherin compared to shSCR control (scale bar = 100μm).

### Loss of PP2A-B56α exacerbates tumorigenic phenotypes in NSCLC cell lines

The process of EMT underlies metastatic phenotypes. Therefore, to determine if the B56α-mediated molecular changes are associated with increased migratory capacity, cells were plated in trans-well migration assay. H1650 shSCR and shB56α cells were plated and allowed to migrate for 24 hours prior to fixing and staining with crystal violet for quantification. Suppression of PP2A-B56α significantly increased the number of migratory cells compared to shSCR control (Figure 3A). To assess if this phenotype is specific to B56α, we transiently overexpressed exogenous B56α (B56αOE) in shB56α cells. Restoration of B56α expression attenuated the migratory capacity of shB56α cells, reducing migration by 30% compared to empty vector control (EV) (Figure 3A, B and Supplemental Figure 2A). To determine the impact of PP2A inhibition on the invasive capacity of H1650 cells, shSCR and shB56α cells were first plated into low attachment round bottom plates to form spheroids and then embedded into media supplemented with 10% Matrigel and imaged over time (96 hours). H1650 shSCR cells formed well organized structures with defined borders and had minimal cell invasion (Figure 3C and Supplemental Figure 2B). In stark contrast, shB56α spheroids were highly invasive with no defined border, as evidenced by an 80% decrease in circularity score where 1 is a perfect circle (Figure 3C, D). Additionally, these structures had an abundance of cells (both single cell and clusters) invade into the surrounding extracellular matrix. Similar to growth in 2D cell culture (Figure 2), these invasive cells expressed high levels of the mesenchymal marker, Vimentin (Supplemental Figure 2C).

**Figure 3:**
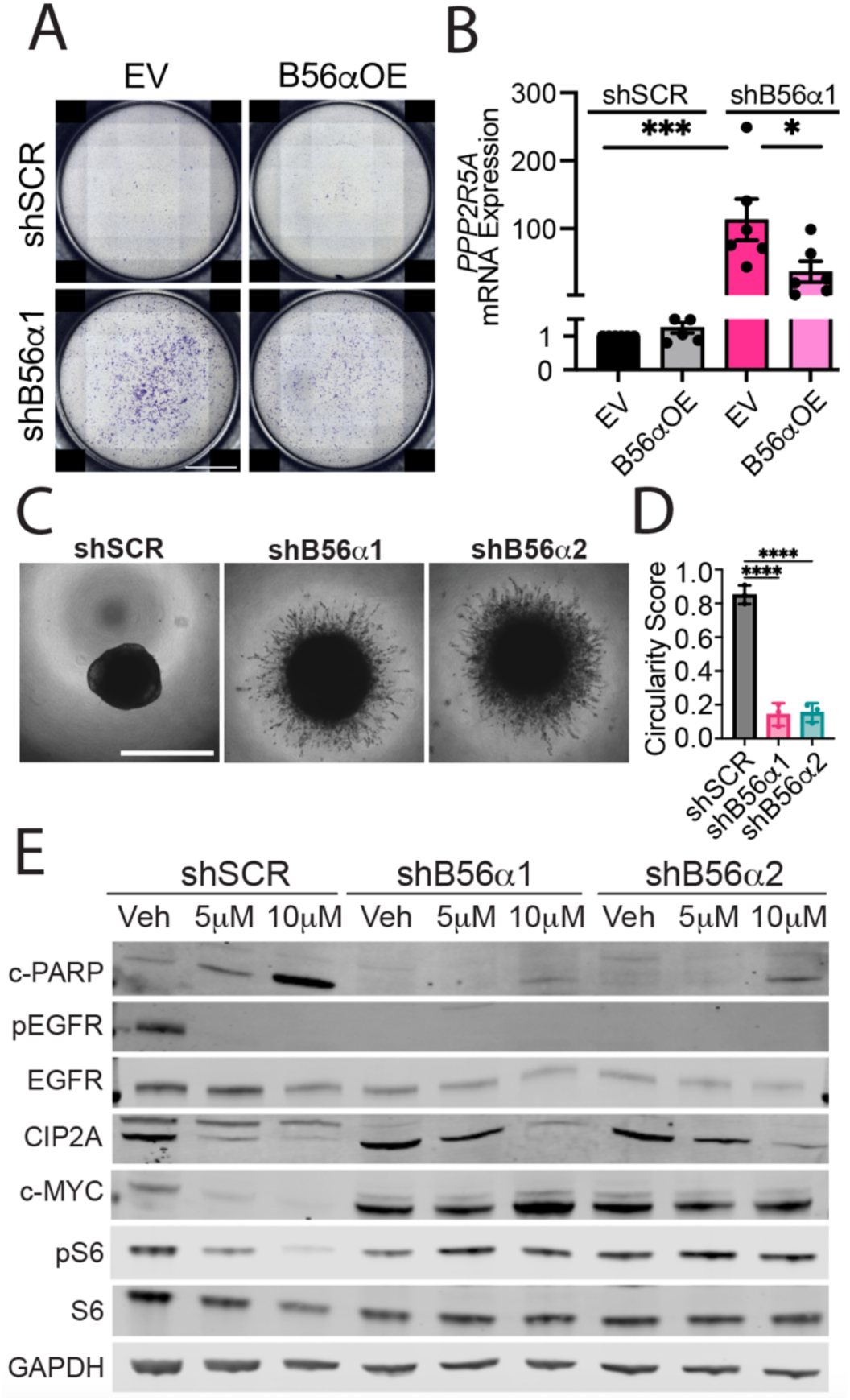
Suppression of B56α increases invasive NSCLC phenotypes. A) Representative full-well images of trans-well migration assay of H1650 KD cells compared to shSCR control, with transient overexpression of B56α (B56αOE) compared to empty vector control (EV). Scale bar = 2mm. B) Quantification of A showing number of cells migrated per well (n=3 biological replicates). B) Representative images of H1650 KD spheroids at 96 hours after embedding in Matrigel (scale bar = 1mm). D) Quantification of D using circularity score. E) Representative western blot images of H1650 KD cells treated with 5μM or 10μM Osimertinib for 24 hours showing cleaved PARP (c-PARP), phospho-EGFR (p-EGFR;Y1068), total EGFR, CIP2A, c-MYC, phospho-S6 (pS6; S240/244), total S6, and GAPDH compared to vehicle (DMSO) control.

To determine if the shift towards a mesenchymal cell state in response to B56α inhibition is associated with a loss of dependency on EGFR signaling, H1650 shB56α cell lines were treated with increasing doses of the EGFR inhibitor Osimertinib for 24 hours and then subjected to Western Blot analysis. Drug doses were chosen based on the published H1650 Osimertinib IC50 of 2.5μM (DepMap) after 72 hrs [26]. In shSCR control cells, Osimertinib treatment reduced EGFR phosphorylation at Tyrosine1068 (pY1068), a marker of EGFR activation, and induced the cleavage of poly (ADP-ribose) polymerase (PARP), a marker of apoptosis (Figure 3E). In contrast, suppression of B56α resulted in an almost complete loss of EGFR pY1068 levels at baseline in vehicle treated conditions; however, the loss of phosphorylation at this site is indirect, as PP2A is a serine/threonine phosphatase. Paradoxically, despite the loss of phospho-EGFR activation, shB56α cells displayed increased proliferation rates (Supplemental Figure 2D), potentially indicating that these cells have become uncoupled from EGFR signaling pathways. Consistent with EGFR-independent survival, the induction of apoptosis (cleaved-PARP) in response to Osimertinib was attenuated in shB56α cells compared to shSCR controls (Figure 3E). Previous studies have determined that cytotoxic responses to EGFR inhibitors are mediated in part through decreased CIP2A expression and increased PP2A activity [27,28]. However, we found that Osimertinib treatment decreased CIP2A expression in all conditions, but was only effective at killing B56α wild type cells. Similarly, PP2A-B56α activation is known to drive the degradation and turnover of MYC protein. In shB56α cells, both MYC expression and a MYC mediated pathway, phospho-S6 Kinase, were dramatically increased in response to B56α knockdown and remained high even in the presence of Osimertinib. These results highlight a crucial role for B56α in regulating invasive NSCLC phenotypes downstream of EGFR activation potentially through the activation of MYC-dependent pathways.

### Overexpression of PP2A-B56α drives an epithelial cell state and suppresses tumor phenotypes

To determine if increased expression of PP2A-B56α shifts cells towards an epithelial cell fate, we analyzed NSCLC cell lines with stable overexpression of PP2A-B56α. HCC827 and H1650 cells were transduced with either empty vector (EV) or HA-tagged B56α to generate stable overexpression lines (B56αOE). Consistent with the dramatic morphological changes that occur with suppression of B56α, H1650 was highly intolerant of B56α overexpression and senesced shortly after transduction (Supplemental Figure 3). PP2A-B56α stable overexpressed in HCC827 resulted in cells shifting to a more epithelial “cobblestone” morphology (Figure 4A, B), which was accompanied by increased expression of E-cadherin as shown by qRT-PCR and western blot (Figure 4C-E). Protein expression of the PP2A-B56α target, MYC, was significantly decreased with B56αOE, indicating that PP2A is active in these cells (Figure 4D, F). Analysis of B56αOE cells by immunofluorescence identified increased E-cadherin positive cell clusters compared to EV control cells. In contrast, there was no significant loss of Vimentin expression, suggesting that B56α overexpression increases epithelial signaling programs (Figure 4G). In addition to increased E-cadherin expression, B56αOE cells were significantly less migratory as shown by a trans-well migration assay compared to EV (Figure 4H, I). Collectively, these findings suggest that low B56α expression increases the propensity of NSCLC cells to undergo cell state changes, while high B56α expression reinforces a more restricted epithelial cell state.

**Figure 4:**
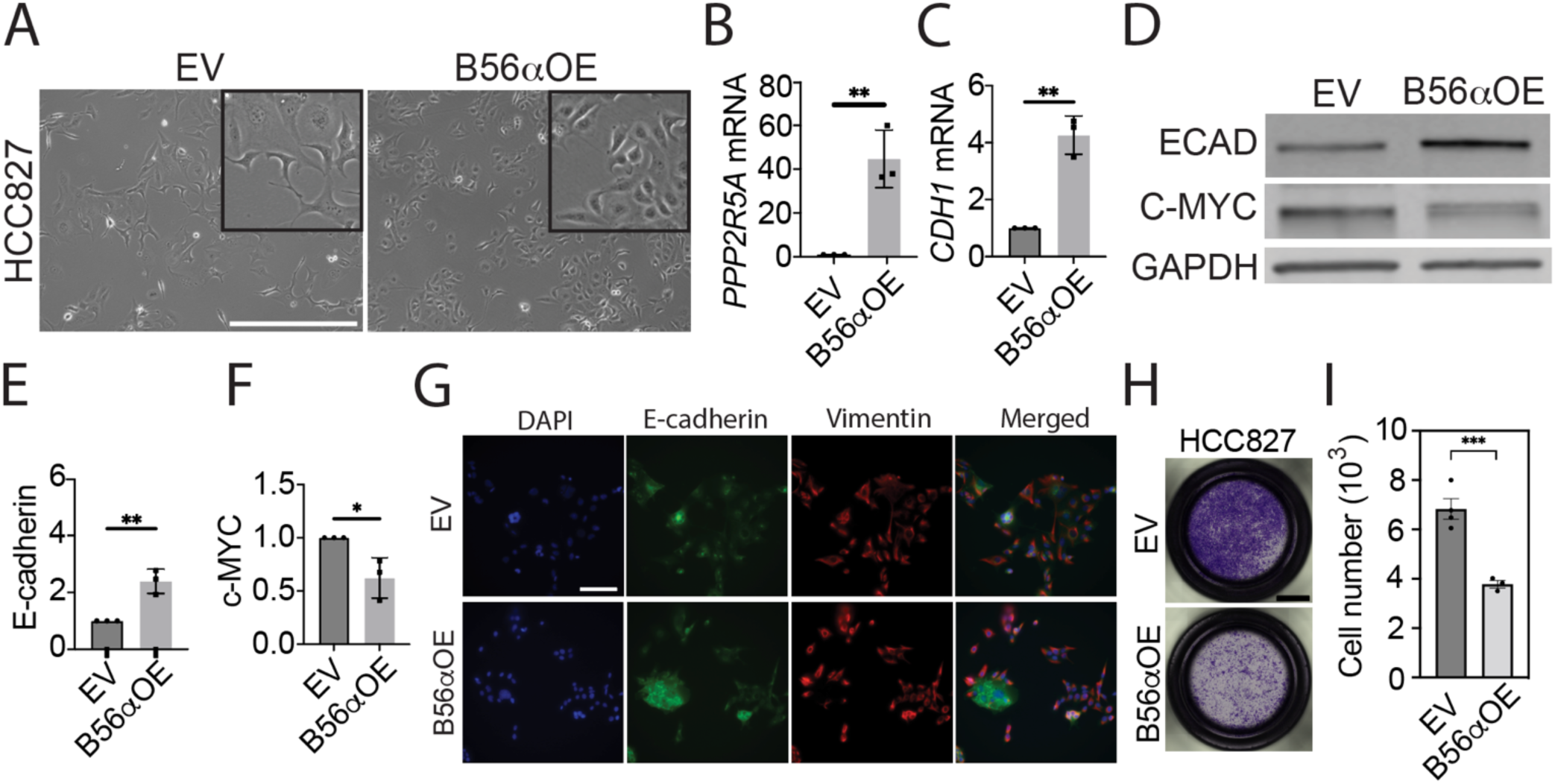
Overexpression of B56α increases E-cadherin expression and reduces migratory capacity. A) Representative brightfield images of HCC827 stable overexpression (B56αOE) compared to empty vector control (EV). Scale bar = 500 μm. B) mRNA expression of B56α (*PPP2R5A*) and C) E-cadherin (*CDH1*) in HCC827 B56αOE cells compared to EV. D) Representative western blot of E-cadherin, c-MYC, and GAPDH protein expression in HCC827 B56αOE compared to EV. E) Quantification of E-cadherin protein and F) c-MYC protein from western blot in panel D (n=3 biological replicates). G) Representative images of HCC827 B56αOE compared to EV showing DAPI, E-cadherin, and Vimentin by immunofluorescent imaging from 3 biological replicates (scale bar = 100μm). H) Representative full-well images of transwell migration assay of HCC827 B56αOE cells compared to EV from 3 biological replicates. I) Quantification of number of cells migrated in panel H (scale bar=2mm).

### Suppression of B56α results in cellular rewiring with large-scale proteomic changes

As knockdown of B56α resulted in dramatic morphological and signaling changes (Figures 2 and 3), we performed global proteomics and phosphoproteomics on H1650 shSCR and shB56α cells to understand the impact of B56α phosphatase inhibition on the NSCLC proteome. H1650 shSCR and shB56α were plated in triplicate and harvested after 48 hours. Proteins were Tandem Mass Tag labeled and analyzed by mass spectrometry using Nano-LC-MS/MS as outlined in Materials and Methods. Proteomic analysis revealed that suppression of B56α resulted in 1,442 significantly changed total proteins, with 785 downregulated and 657 upregulated (Figure 5A and Supplemental Data Table 1). Consistent with a loss of epithelial characteristics, Gene Ontology (GO) enrichment identified several cellular adhesion and cell junction signaling programs as being significantly downregulated in shB56α cells compared to shSCR controls (Figure 5B and Supplemental Data Table 2). Analysis of transcription factor target interactions using TRRUST [29] revealed a decrease in ZEB1 regulated targets in shB56α cells (Supplemental Figure 4A).

**Figure 5:**
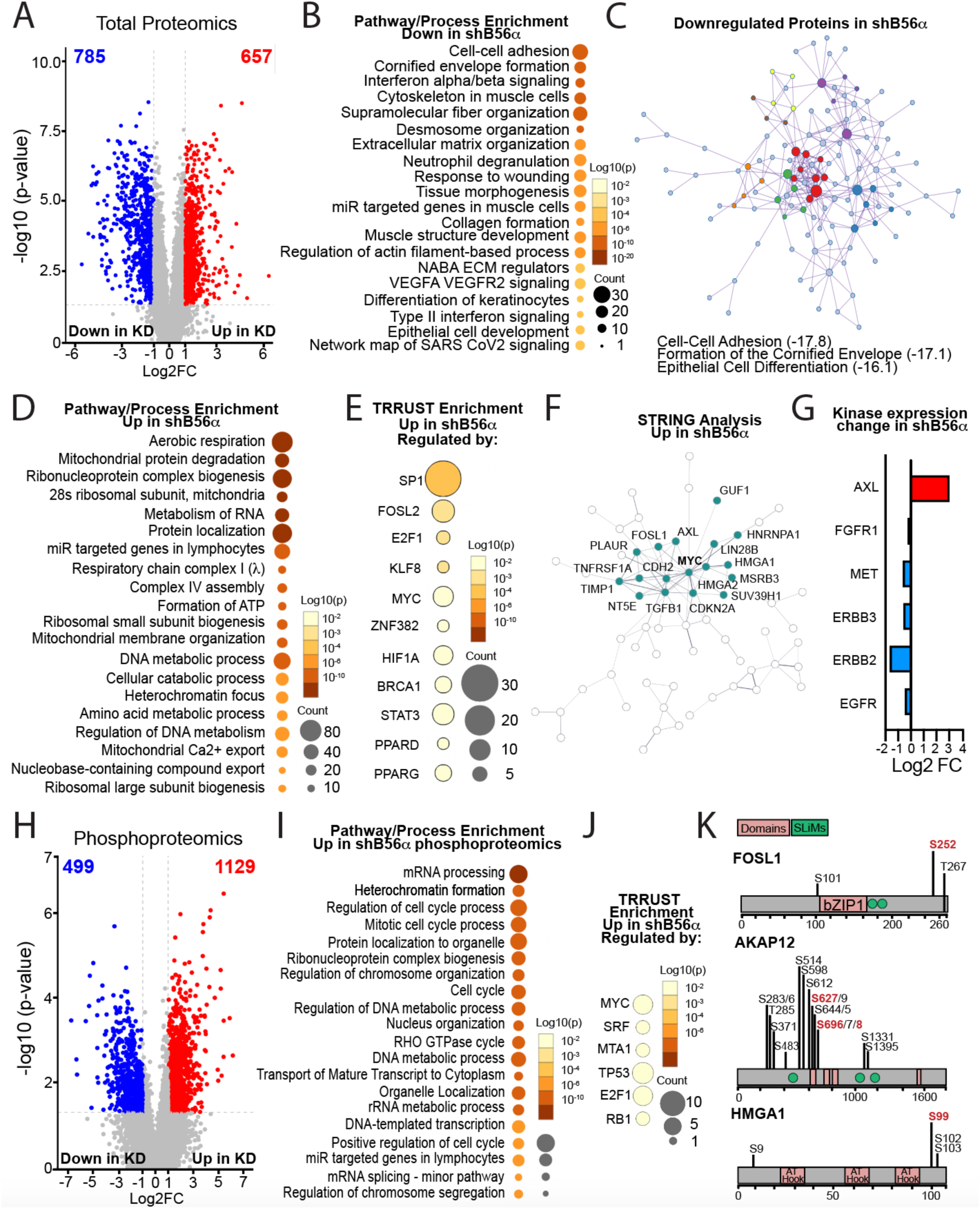
B56α knockdown alters the expression of a diverse set of proteins involved in tumorigenic plasticity. A) volcano plot of total proteomic changes in H1650 KD cells compared to shSCR control (Log_2_FC <-1 or >1; p<0.05) B) Pathway and Process Enrichment Analysis using Metascape analysis of significantly downregulated proteins in total proteomics (Log_2_FC <-2, p<0.05). C) MCODE Metascape analysis of pathway enrichment with protein interactions of significantly downregulated proteins in total proteomics (Log_2_FC <-2, p<0.05). Top 3 enrichment pathways listed with Log_2_ p-value in parentheses. D) Metascape Pathway and Process Enrichment Analysis of significantly upregulated proteins in total proteomics (Log_2_FC >1, p<0.05). E) TRRUST enrichment analysis using Metascape identifying gene sets that are enriched in shB56α and regulated by specific transcription factors. F) STRING analysis of top 15% upregulated proteins in shB56α, highlighting the enriched c-MYC-centered network. G) Curated list of kinases significantly changed in shB56α total proteomics. H) Volcano plot of phosphoproteomic changes in H1650 shB56α (Log_2_FC <-1 or <1; p<0.05). I) Metascape Pathway and Process Enrichment Analysis of significantly upregulated proteins in phosphoproteomics (Log_2_FC >2, p<0.05). J) TRRUST enrichment analysis using Metascape identifying gene sets that are enriched in shB56α phosphoproteomics and regulated by specific transcription factors. K) Significantly upregulated phospho-sites on FOSL1, AKAP12, and HMGA1 in shB56α cells are denoted in black. Red sites indicate known phosphorylation events implicated in EMT or invasion. Major domains (pink). B56 binding motifs, SLiMs (green).

Given these results, we quantified the expression of classical EMT-TFs including *ZEB1*, SNAIL (*SNAI1*), SLUG (*SNAI2*), and TWIST (*TWIST1*). ZEB1 mRNA expression was increased in shB56α cells, albeit in a highly variable manner, whereas *SNAI1*, *SNAI2*, and *TWIST* mRNA expression were decreased comparatively (Supplemental Figure 4B). These results suggest that the cellular plasticity that occurs in response to PP2A-B56α suppression most likely does not function through classical EMT-TF signaling and may represent a unique posttranslational mechanism. Using the Metascape Molecular Complex Detection (MCODE) [30] algorithm to identify significant direct protein-protein interaction networks, we determined that cell-cell adhesion, formation of the cornified envelope, and epithelial cell differentiation networks were significantly enriched in the downregulated proteins (Figure 5C and Supplemental Data Table 3). Within individual MCODE clusters (MCODE 1-7) there was significant enrichment in desmosome, actin, intermediate filament, and E-Cadherin organization (Supplemental Figure 4C and Supplemental Data Table 3). These findings, taken together with the fact that the majority of significantly altered proteins were downregulated in shB56α cells (785 versus 657), suggest that PP2A-B56α may function to maintain NSCLC epithelial differentiation programs in order to suppress cellular plasticity.

Within the significantly upregulated proteins in shB56α cells, there was a strong enrichment for signaling programs regulating cell cycle, as well as chromatin organization and mRNA processing (Figure 5D and Supplemental Data Table 4). These enrichments were consistent with the increased proliferative capacity of shB56α cells (Supplemental Figure 2D) and support a potential role for B56α in regulating large-scale gene expression programs. Indeed, similar enrichments have been previously identified in response to either a loss of the PP2A catalytic or A subunits [31]. Analysis of transcription factor target interactions identified an enrichment of genes regulated by transcription factors implicated in oncogenesis and EMT, including SP1, FOSL2, KLF8, and MYC (Figure 5E and Supplemental Data Table 5)[32–40]. Notably, classical EMT-TFs such as ZEB1 were not enriched in shB56α cells, supporting the use of alternative pathways to drive EMT. Additionally, there was a strong MYC-centered network found within the top 15% of upregulated proteins that was significantly enriched for factors that regulate EMT signaling programs, including Transforming growth factor beta 1 (TGFB1), Tissue Inhibitor of Metalloproteinases 1 (TIMP1), N-cadherin (CDH2), and High Mobility Group AT-Hook 2 (HMGA2) (Figure 5F and Supplemental Figure 4D). Of particular interest, this node included angiotoxin receptor-like (AXL), a driver of EMT and EGFRi therapeutic resistance in NSCLC [4,41]. Furthermore, of the kinases implicated in EGFR mutant NSCLC tumor progression, AXL was the only factor upregulated with B56α suppression (Figure 5G) and is known to increase MYC activity to exacerbate oncogenesis, potentially implicating a broad regulation of these convergent pathways [4,41,42]. Finally, consistent with studies supporting a role for PP2A in epigenetics [31,43], there were 71 significantly altered proteins involved in the epigenetic regulation of gene transcription (GO: 0040029), most likely contributing in part to the large-scale protein expression changes seen in shB56α cells.

Consistent with the phosphatase function of PP2A-B56α, there were 1,129 phospho-sites found to be significantly increased in shB56α cells, compared to 499 that were significantly downregulated (Log_2_ >1, *p<*0.05) (Figure 5H and Supplemental Data Table 6). The unique proteins that displayed increased phosphorylation in shB56α cells were enriched for cell cycle, RNA splicing and processing, and transcriptional regulation pathways, as well as MYC regulated target genes, (Figure 5I, J and Supplemental Data Table 7 and 8). As a validation of our system, we were able to identify at least 20 phospho-sites implicated to be regulated by PP2A (Supplemental Data Table 9). Furthermore, of the top 100 upregulated phospho-proteins, 70% contained a predicted PP2A-B56 specific short linear binding motifs (SLiMs) [44], indicating that both direct and indirect PP2A targets are altered in shB56α cells. Within the top 100 differential phosphorylated proteins, there were several upregulated phospho-sites on factors strongly implicated in NSCLC metastasis or EMT, such as Vimentin (a known PP2A regulated target), Fos-like antigen 1 (FOSL1), High Mobility Group AT-Hook 1 (HGMA1), and A-kinase anchoring protein 12 (AKAP12) (Figure 5K and Supplemental Figure 4E) [45–47]. Combined, the changes in the H1650 phosphoproteome suggest that suppression of B56α dynamically rewires cells, potentially alleviating the tight regulation of pathways necessary to maintain cellular homeostasis and, thereby, reducing the threshold for cellular plasticity.

### Suppression of PP2A-B56α enhances the metastatic potential of NSCLC *in vivo*

EMT has been associated with increased metastatic capability in a wide variety of tumor types [48]. Our findings suggest that suppression of PP2A-B56α results in EMT *in vitro*. Therefore, to determine the impact of B56α signaling on metastasis *in vivo*, 2 million H1650 shSCR and shB56α cells were injected intravenously into immune-compromised mice and tumors were allowed to develop over six weeks, at which point lung and liver tissues were harvested for immunohistological analysis. The majority of shB56α lungs had large, macroscopic tumor nodules and/or widespread tumor dissemination compared shSCR as seen by both gross morphology and histology (H&E) (Figure 6A). To quantify lung colonization, immunofluorescence for the human marker ATP-dependent DNA helicase II subunit 2 (Ku-80) was performed and quantified using QuPath [49]. There was a significant ∼5-fold increase in the percent of Ku-80+ tumor cells within shB56α lungs compared to shSCR controls (Figure 6A, B). In addition to lung tumors, almost all mice injected with shB56α cells formed large macroscopic liver tumors compared to mice injected with shSCR cells, which had almost no liver outgrowth (Figure 6C and Supplemental Figure 5A). Furthermore, in many cases the shB56α tumor tissue was so extensive that almost no normal liver tissue remained (Supplemental Figure 5B). Similar to our *in vitro* results, the Ku-80+ shSCR tumor cells that were able to colonize the lung remained small clusters that were predominately E-cadherin positive and Vimentin negative (Figure 6D). In contrast, shB56α tumors remained E-cadherin negative and Vimentin positive, indicative of an aggressive EMT-like state. These results support our hypothesis that PP2A-B56α functions to suppress NSCLC cellular plasticity and suggest that inhibition of B56α may contribute to both EMT and metastatic potential.

**Figure 6:**
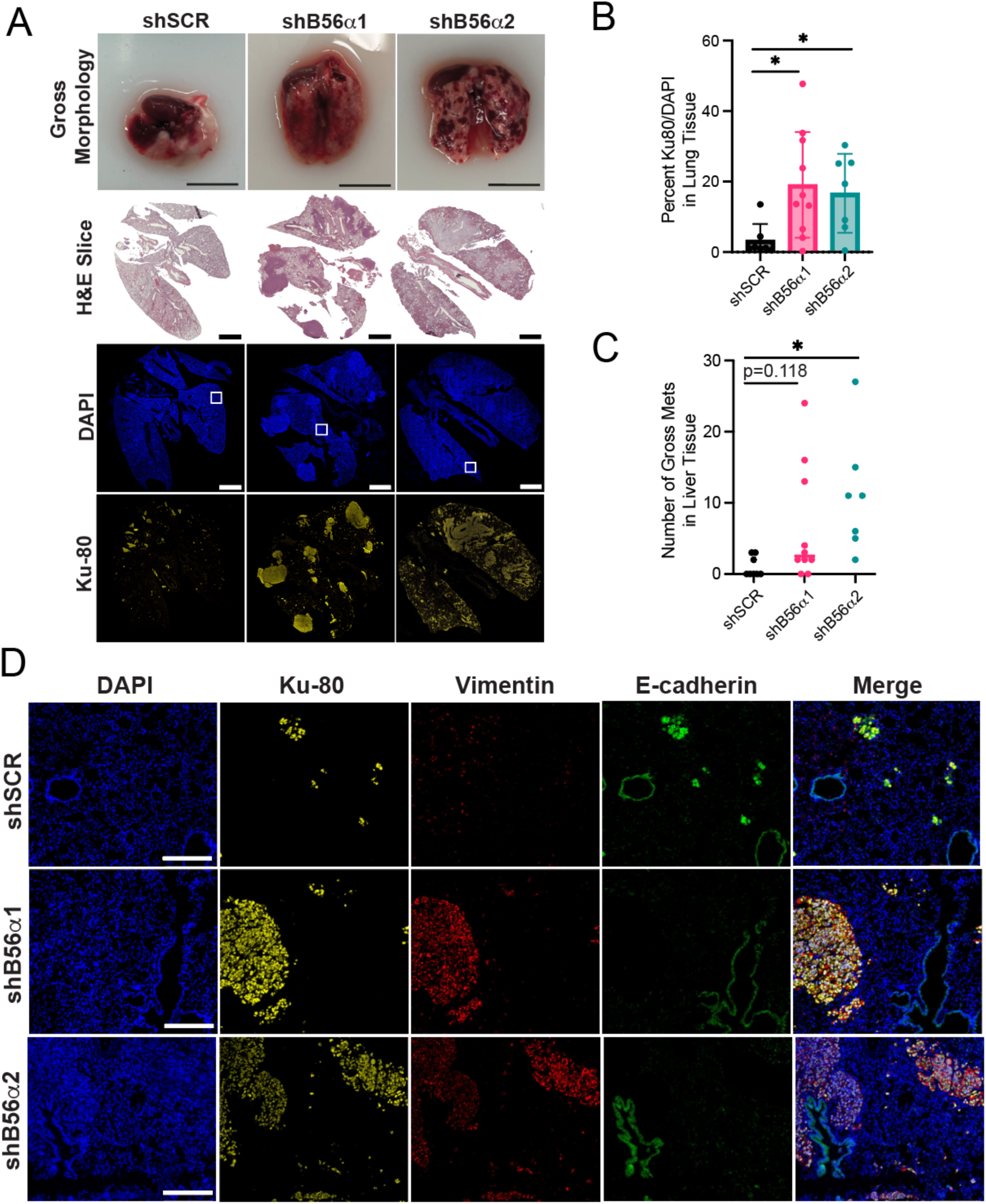
B56α negatively regulates NSCLC metastatic potential *in vivo*. A) Representative gross morphology images (scale bar = 1cm), H&E slices (scale bar = 2mm), and immunofluorescence for DAPI and Ku-80 (scale bar = 2mm) in lungs harvested from mice 6 weeks post-tail vein injection with either shB56α or shSCR cells. B) Quantification of the percent of Ku-80 positive nuclei in the lung from shB56α and shSCR injected mice at endpoint. C) Quantification of the number of gross liver tumor nodules in each condition. For panel B and C, Brown-Forsythe and Welch ANOVA, *p<0.05. D) Representative immunofluorescence images of DAPI, Ku-80, Vimentin, and E-cadherin taken from the regions denoted panel A (white boxes) (scale bar = 250μm).

## DISCUSSION

PP2A functions as key gatekeeper to phosphorylation cascades and provides a key “off” switch to a large number of pathways implicated in cancer. However, our understanding of how PP2A dysregulation impacts NSCLC cellular plasticity and invasion is poorly understood. We have determined that suppression of the specific PP2A subunit, B56α, allows NSCLC cells to lose their epithelial characteristics and gain mesenchymal characteristics, indicative of an EMT. Furthermore, identified changes in EMT markers were accompanied by a significant increase in migratory and invasive capacity as evidenced by 2D migration and 3D spheroid invasion assays *in vitro*, and tail vein invasion assay *in vivo*. These invasive phenotypes were associated with a vast number of significant proteomic and phosphoproteomic changes within proteins implicated in invasion, EMT, proliferation, and transcriptional regulation. Together, these findings indicate that PP2A-B56α functions as a key modulator of cell state in NSCLC and that suppression of this subunit leads to drastic cellular reprogramming that leads to aggressive disease.

B56α is only one of many PP2A B subunits (4 families, 16 subunits total in addition to splice variants). Within the four B subunit families there is a high degree of structural homology suggesting potential functional redundancy [50]. Yet, the suppression of B56α alone leads to drastic morphological and molecular changes in the EGFR mutant NSCLC cell lines tested. Conversely, the overexpression of B56α increases epithelial markers and suppresses invasive phenotypes. These results indicate that the regulation of NSCLC EMT phenotypes is likely to be B56α specific. However, we cannot rule out the possibility that suppression B56α alters the dynamics of the PP2A holoenzyme composition allowing for other subunits to partially contribute to the observed phenotypes. The spatial and temporal relationships that occur between PP2A subunits are both highly biologically relevant and technically challenging to answer; therefore, this area of research remains an open question within the PP2A field. Given that a large percent of the top differentially regulated proteins contain B56-specific SLiM domains, it is also possible that many of these targets are co-regulated by multiple PP2A B56 subunits. There are limited examples of such regulation, for example B<u>55</u>α and B<u>56</u>α display opposing roles in the regulation of MYC at two different phospho-sites in breast cancer [51]. However, since the function of individual PP2A B subunits is highly dependent on cellular context, further studies are necessary to identify direct B56α targets versus PP2A as a whole. Finally, as small molecule activators of PP2A (SMAPs) are currently being tested in preclinical studies [18,21,50], our findings suggest that pharmacological activation of PP2A may be efficacious in metastatic EGFR-driven NSCLC. In fact, previous studies have demonstrated that SMAPs reduce subcutaneous xenograft tumor growth in EGFR tyrosine kinase inhibitors (TKI) resistant NSCLC [52].

Through our phospho-proteomics analysis, we identified a significant downregulation of proteins involved in cell-cell adhesion and an upregulation of factors involved in EMT, including both MYC and MYC regulated pathways. B56α is known to dephosphorylate MYC at S62 resulting in MYC’s proteasomal degradation [18,20,21,23]. Therefore, enrichment of MYC signaling pathways highlights the strong reliance of EGFR mutant NSCLC on the posttranslational regulation of this transcription factor by PP2A and implicates disruption of the PP2A-MYC axis as a potential driver of EMT. In addition to MYC there were a wide variety of factors implicated in the mesenchymal reprogramming of cancer cells including HMGA1/2, FOSL1, and TGFβ. As a whole, EMT is a process driven by transcriptional reprogramming in response to increased expression and/or activation of classical EMT-TF such as ZEB1, SNAI1, and TWIST1. In shB56α cells, ZEB1 mRNA was increased but ZEB1 transcriptional targets were negatively enriched at the protein level. These findings suggest that EMT in response to B56α suppression likely does not function through classical EMT-TFs. Instead, the disruption of PP2A’s posttranslational regulation may represent an alternative path towards EMT. Taken together with the loss of epithelial markers, we hypothesize that the B56α functions to restrict posttranslational signaling in order to maintain epithelial cell states. The suppression of this subunit potentially alleviates this restrictive programming poising cells for EMT. This hypothesis is further supported by our finding that expression of exogenous B56α for just 48 hours is able to significantly reduce migratory phenotypes in shB56α cells, indicating that the posttranslational regulation of EMT by PP2A is highly transient and dynamic in nature.

Given the significant impact of EGFR signaling on NSCLC progression, therapeutic strategies to target this pathway have provided immense clinical benefit, with EGFR TKIs such as Osimertinib resulting in advanced NSCLC patient survival increasing from 4 months to over 10 months when compared to platinum-based therapies. However, in a large portion of patients (45%), resistance to these targeted therapies occurs in less than a year [53]. Suppression of the B56α subunit of PP2A leads to a near complete loss of Y1068 p-EGFR, suggesting that shB56α cells function independently of EGFR signaling, despite mutant EGFR (E19 del) being the driving mutation in these cells. EMT drives therapeutic resistance in a broad range of tumors and therapeutics, including EGFRi in NSCLC. While our studies suggest that shB56α have reduced Osimertinib efficacy, future studies will aim to interrogate the dependency of PP2A-regulated EMT on therapeutic response, whether PP2A perturbation in primary patient tumors (e.g. high CIP2A (PP2A inhibitor)) alters therapeutic response.

In summary, these studies address a critical gap in our understanding of the posttranslational mechanisms that govern NSCLC EMT. Our findings strongly implicate PP2A-B56α as a critical regulator of cellular plasticity in EGFR mutant NSCLC and highlight the need for continued studies interrogating the therapeutic potential of PP2A in the metastatic setting.

## MATERIALS AND METHODS

### Cell Culture

All primary and established NSCLC cell lines were cultured in RPMI-1640 (Fisher SH3002701) containing 10% fetal bovine serum (Fisher FB12999102). Primary cell lines were previously established from patients following core biopsy or pleural effusion as previously described [24,25]. Patients signed informed consent to give permission for research to be completed using their samples through a Dana-Farber-Harvard Cancer Center Institutional Review Board approved protocol. To generate B56α overexpression cells, B56α was cloned from the pCEP-4HA B56alpha plasmid (Addgene #14532) into pSIN vector [54]. Lentivirus was then used to establish stable HA-tagged B56α overexpression and shRNA B56α knockdown cell lines. Lentivirus was produced using HEK293T cells transfected using Lipofectamine 3000 (Fisher Scientific, L3000015), the plasmid of interest, and packaging plasmids (pAX.2 Addgene Plasmid #35002 and pMD2.G Addgene Plasmid #12259). Parental cell lines were transduced with viral media containing the plasmid of interest (pSIN or SMARTvector shRNA plasmids; VSC11709 (shSCR), V3SH11240-225202507 (shB56α1), V3SH11240-229314505 (shB56α2), Horizon Discovery) and selected with the appropriate antibiotic. All cell lines were routinely tested for Mycoplasma using PCR-based strategies and grown at 37C in 5% CO_2_ atmosphere. For B56α rescue experiments, cells were seeded and transiently transfected with HA-tagged B56α using Lipofectamine 3000 (Fisher Scientific, L3000015) according to manufacturer instructions.

### Quantitative RT-PCR

RNA was isolated using the ThermoScientific GeneJet RNA Purification Kit (Thermofisher, #K0732). cDNA was generated using the High-Capacity cDNA Reverse Transcription Kit (Fisher Scientific, #43-688-14). Quantitative RT-PCR was performed using PowerUp SYBR Green Master Mix reagent (Fisher Scientific, #A25743) on the QuantStudio 3 with the indicated primers (Supplemental Methods Table 1). The fold change relative to vehicle/control was analyzed using the 1ΔΔ(C_t_) method.

### Western blotting

Cells were lysed (20mM Tris pH7.5, 50mM NaCl, 0.5% Triton X-100, 0.5% deoxycholate, 0.5% SDS, and 1mM EDTA) and protein concentration was determined using the DC protein assay kit (Bio-Rad, 5000112). Lysate was prepared using XT Sample Buffer (Bio-Rad, 1610791) and XT Reducing Agent (Bio-Rad, 1610792) then boiled at 95°C. SDS-PAGE was run on 4-12% gradient bis-tris protein gel (Bio-Rad, 3450123) and transferred to PVDF membrane. Membrane was blocked in Licor TBS blocking buffer (Fisher Scientific, NC1660550) for 1 hour at room temperature and incubated with primary antibody overnight at 4°C (Supplemental Methods Table 2) and then incubated with secondary antibody for 1 hour at room temperature. Membranes were scanned using the Licor Odyssey DLx Imaging System and analyzed using Image Studio software v5.2.5. For drug treatment studies, cells were seeded in 6-well plates and Osimertinib (Medchem 1421373-65-0, Lot 273365) was immediately added to the culture upon plating. After 24 hours, cells were lysed and processed for western blot analysis.

### Immunocytochemistry

Cells were seeded on top of sterile glass coverslips in 24-well cell culture plates. After 48 hours, cells were fixed in 4% paraformaldehyde for 20 minutes at room temperature. Cells were permeabilized by 40μg/mL digitonin (Sigma Aldrich, D141-100MG) for 1 minute. Slides were incubated at 4°C overnight in primary antibody solutions (diluted with 2% bovine serum albumin (BSA) in PBS). Slides were incubated in secondary antibody solutions (diluted in 2% BSA in PBS) for 1 hour at room temperature and DAPI stained for 5 minutes at room temperature (Supplemental Methods Table 3). Coverslips were added to slides using ProLong Gold anti-fade reagent (Invitrogen, P36934). All rinses between solutions were done with PBS-T (phosphate buffered saline and 0.05% Tween-20). Slides were imaged with Nikon Ni-U Upright Microscope with epifluorescence and processed with FIJI software v2.14.0.

### Transwell Migration Assay

Transwell inserts were soaked in serum-free RPMI-1640 media for one hour prior to seeding cells. Cells were seeded at a density of 2×10^4^ per well in serum-free RPMI-1640 with RPMI-1640 containing 10% FBS in the lower chamber. After incubating for 48 hours, or 24 hours for rescue experiments, the media was aspirated and stained with crystal violet for 1.5 hours. Migrated cells were imaged with the EVOS M7000 and quantified with FIJI software.

### Spheroid Invasion Assay

50,000 cells per well were plated in 96-Well Clear Ultra Low Attachment Microplates (Corning, 07-201-680). After 72 hours, spheroids were removed from low attachment plates and embedded by plating on top of 80% Matrigel (Corning 356231, lot: 2024001) and suspended in 10% Matrigel. Spheroids were imaged every 24 hours for 96 hours at which point spheroids were fixed and processed for immunofluorescent imaging. Spheroid circularity was calculated using FIJI software v2.14.0.

### Proliferation Assays

Proliferation: 100,000 cells were plated in triplicate in 4, 6-well plates. Cell number was quantified every 24 hours. Population doubling: 100,000 cells were plated in triplicate into 6-well plates. Every 72 hours, cells were trypsinized and cell number was quantified. Cells were then passaged by replating 100,000 cells onto a new plate. Cells were quantified over a total of 12 days and population doubling was calculated using natural log.

### Phosphoproteomics

Sample preparation, mass spectrometry analysis, bioinformatics, and data evaluation for quantitative proteomics experiments were performed in collaboration with the Indiana University Proteomics Center for Proteome Analysis at the Indiana University School of Medicine similarly to previously published protocols [55,56].

#### Protein Extraction and Digestion

Cell pellets were resuspended in 100 µL 8 M Urea, 100 mM Tris pH 8.5. The resuspended cell pellets were transferred to Diagenode Bioruptor tubes (Cat No: C30010010). Cells were lysed via Diagenode Bioruptor, 30s on/30s off, for 30 cycles. Samples were then clarified by centrifuging for 30 min at 12,000 rcf. Supernatants were analyzed in a Bradford assay (Biorad Cat No: 5000002) to determine protein concentration. 1 mg of each sample was treated with 5 mM tris (2-carboxyethyl) phosphine hydrochloride (Sigma-Aldrich Cat No: C4706) to reduce disulfide bonds and the resulting free cysteine thiols were alkylated with 10 mM chloroacetamide (Sigma Aldrich Cat No: C0267). Samples were diluted with 50 mM Tris.HCl pH 8.5 (Sigma-Aldrich Cat No: 10812846001) to a final urea concentration of 2 M for overnight Trypsin/Lys-C digestion at 35 °C (1:25 protease:substrate ratio, Mass Spectrometry grade, Promega Corporation, Cat No: V5072).

#### Peptide Purification and phosphopeptide enrichment

Digestions were quenched with trifluoroacetic acid (TFA, 0.5% v/v) and desalted on Waters Sep-Pak® Vac cartridges (Waters™ Cat No: WAT054955) with a wash of 1 mL 0.1% TFA followed by elution in 3x 0.2 mL of 70% acetonitrile 0.1% formic acid (FA). Peptides were dried by speed vacuum. Samples were resuspended in phosphopeptide binding buffer and phosphopeptides were enriched using Thermo Fisher Scientific High Select TiO2 tips (Cat No A32993) according to manufacturer’s instructions. Flow through (non-phosphopeptides) and phosphopeptides were dried down by speed vacuum.

#### TMTpro labeling

An equivalent of 50 μg of the global (non-phospho) peptides and all of each phosphopeptide enrichment were resuspended in 100 mM triethylammonium bicarbonate (TEAB, pH 8.5 from 1 M stock). Each sample was then labeled overnight at room temperature, with 0.5 mg of Tandem Mass Tag Pro (TMTpro™) reagent (16-plex kit, manufactures instructions Thermo Fisher Scientific, TMTpro™ Isobaric Label Reagent Set; Cat No: 44520, lot no. ZA382395 see Table X below). Reactions were quenched with 0.3 % hydroxylamine (v/v) at room temperature for 15 minutes. Labeled peptides were then mixed and dried by speed vacuum.

#### High pH Basic Fractionation

Half of the combined global sample and all of the phosphopeptide sample were resuspended in 0.5% TFA and fractionated on a Waters Sep-Pak® Vac cartridge (Waters™ Cat No: WAT054955) with a 1 mL wash of water, 1 mL wash of 5% acetonitrile, 0.1% triethylamine (TEA) followed by elution for the global sample in 8 fractions of 12.5%, 15%, 17.5%, 20%, 22.5%, 25%, 30%, and 70% acetonitrile, all with 0.1% TEA).

#### Nano-LC-MS/MS

Mass spectrometry was performed utilizing an EASY-nLC 1200 HPLC system (SCR: 014993, Thermo Fisher Scientific) coupled to Eclipse™ mass spectrometer with FAIMSpro interface (Thermo Fisher Scientific). Each multiplex was run on a 25 cm Aurora Ultimate TS column (Ion Opticks Cat No: AUR3-25075C18) in a 50 °C column oven with a 180-minute gradient. For each fraction, 2% of the sample was loaded and run at 350 nl/min with a gradient of 8-38%B over 98 minutes; 30-80% B over 10 mins; held at 80% for 2 minutes; and dropping from 80-4% B over the final 5 min (Mobile phases A: 0.1% formic acid (FA), water; B: 0.1% FA, 80% Acetonitrile (Thermo Fisher Scientific Cat No: LS122500)). The mass spectrometer was operated in positive ion mode, default charge state of 2, advanced peak determination on, and lock mass of 445.12003. Three FAIMS CVs were utilized (-45 CV; -55 CV; -65CV) each with a cycle time of 1 s and with identical MS and MS2 parameters. Precursor scans (m/z 400-1600) were done with an orbitrap resolution of 120000, RF lens% 30, 50 ms maximum inject time, standard automatic gain control (AGC) target, minimum MS2 intensity threshold of 2.5e4, MIPS mode to peptide, including charges of 2 to 6 for fragmentation with 60 sec dynamic exclusion shared across the cycles excluding isotopes. MS2 scans were performed with a quadrupole isolation window of 0.7 m/z, 34% HCD collision energy, 50000 resolution, 200% AGC target, dynamic maximum IT, fixed first mass of 100 m/z.

#### Mass spectrometry Data Analysis

Resulting RAW files were analyzed in Proteome Discover™ 2.5.0.400 (Thermo Fisher Scientific) [57] with a *Mus musculus* UniProt reference proteome FASTA (downloaded 051322) plus common laboratory contaminants (73 sequences). SEQUEST HT searches were conducted with full trypsin digest, 3 maximum number missed cleavages; precursor mass tolerance of 10 ppm; and a fragment mass tolerance of 0.02 Da. Static modifications used for the search were: 1) TMTpro label on peptide N-termini 2) TMTpro label on lysine (K) and 3) carbamidomethylation on cysteine (C) residues. Dynamic modifications used for the search were 1) oxidation on M (2) phosphorylation on S, T or Y, 3) acetylation on protein N-termini, 4) methionine loss on protein N-termini or 5) acetylation with methionine loss on protein N-termini. A maximum of 3 dynamic modifications were allowed per peptide. Percolator False Discovery Rate was set to a strict setting of 0.01 and a relaxed setting of 0.05. IMP-ptm-RS node was used for all modification site localization scores. Values from both unique and razor peptides were used for quantification. In the consensus workflows, peptides were normalized by total peptide amount with no scaling. Unique and razor peptides were used and all peptides were used for protein normalization and roll-up. Quantification methods utilized TMTpro isotopic impurity levels available from Thermo Fisher Scientific. Reporter ion quantification filters were set to an average S/N threshold of 5 and co-isolation threshold of 30%. Resulting grouped abundance values for each sample type, abundance ratio values; and respective p-values (Protein Abundance based with ANOVA individual protein based) from Proteome Discover were exported to Microsoft Excel. Pathway enrichment was performed using the online tool Metascape v3.5 (http://metascape.org) [58].

### Tail Vein Injection Mouse Study

All animal studies were completed in compliance with Purdue University (West Lafayette, IN) animal care and use guidelines after approval by the Purdue Institutional Animal Care and Use Committee. NOD.Cg-Rag1^tm1Mom^ Il2rg^tm1Wjl^/SzJ (NRG) mice (JAX 007799) were used for the tail vein injection study. Mice were injected intravenously with 2 million H1650 cells in 100μL PBS. At 6 weeks post-injection, mice were euthanized by CO_2_ followed by cervical dislocation. Lung and liver were removed from each mouse, perfused and inflated with PBS/heparin, and then fixed in 10% neutral buffered formalin and paraffin embedded.

### H&E Tissue Staining and Immunofluorescence (IF)

For both H&E and IF, lung and liver tissues were sectioned at 6μm thickness using a Thermo HM355S microtome. Slides were baked overnight at 55 °C and H&E staining was performed as previously described [18]. For IF, slides were rehydrated and double antigen retrieval was used (high pH cat:00-4956-58; low pH cat:00-4955-58). Slides were blocked with 3% BSA in PBS for 1 hour at room temperature. Primary antibodies were used as indicated and incubated overnight at 4 °C (Supplemental Table 3). Tissues were washed with PBS-T and incubated in secondary antibodies for 1 hour and DAPI stained for 5 minutes at room temperature. Cells were mounted with Prolong Gold mounting medium and imaged on a Nikon Ni-U Upright Microscope.

### Statistical Considerations

All experiments were performed in at least three independent biological replicates and graphs are plotted with standard deviation (s.d.) unless otherwise denoted in the figure legend. Statistical significance was determined using a two-tail student’s t-test (2 samples) or a one-way ANOVA with post hoc analysis using GraphPad Prism (GraphPad Software)(3 samples or more). If experimental results displayed unequal variance by Bartlett’s test, then a Brown-Forsythe and Welch’s one-way ANOVA test was performed. Values less that p < 0.05 were considered significant. *\*p<0.05, **p<0.01, ***p<0.001, ****p<0.0001*

## ACKNOWLEDGMENTS

We would like to thank Dr. Andrea Kasinski (Purdue University, West Lafayette, IN) for providing established NSCLC cell lines and Dr. Aaron Hata (Harvard University, Massachusetts General Hospital, Boston, MA) for providing the primary patient cell lines. We would like to thank the Purdue Institute for Cancer Research (NIH grant P30 CA023168) and the Purdue Histology Core for their contributions to the data produced in this publication. We would also like to thank all the members of the BAP lab for editing of the manuscript and providing constructive feedback. The mass spectrometry work performed in this work was done by the Indiana University School of Medicine Proteomics Core. Acquisition of the IUSM Proteomics core instrumentation used for this project was provided by the Indiana University Precision Health Initiative. The proteomics work was supported, in part, by the Indiana Clinical and Translational Sciences Institute (funded in part by Award Number UL1TR002529 from the National Institutes of Health, National Center for Advancing Translational Sciences, Clinical and Transltional Sciences Award) and, in part, by the IU Simon Comprehensive Cancer Center Support Grant (Award Number P30CA082709 from the National Cancer Institute). Additional funding for these studies includes: Purdue Institute for Cancer Research SIRG Graduate Research Assistantship (BNH), the Ralph W. and Grace M. Showalter Research Trust (BLA-P), and NIH R01 CA137008 (ANH).

## AUTHOR CONTRIBUTIONS

BLA-P and BNH designed experiments. BNH, GB, CMP, MBB, AKD, SJC, NLA, MRO, and BLA-P performed and analyzed experiments. EHD, WS-K, and KH performed Mass spec sample preparation and EHD performed LC-MS/MS, Data Analysis, Evaluation, Verification, and Methods Write-Up. ANH provided key resources and expertise. BLA-P secured funds and provided supervision. BLA-P and BNH wrote and revised the manuscript. All authors reviewed and finalized the manuscript.

