## Supplemental Figure 1 for "Suppression of PP2A-B56α Drives EMT in EGFR Mutant Non-Small Cell Lung Cancer"

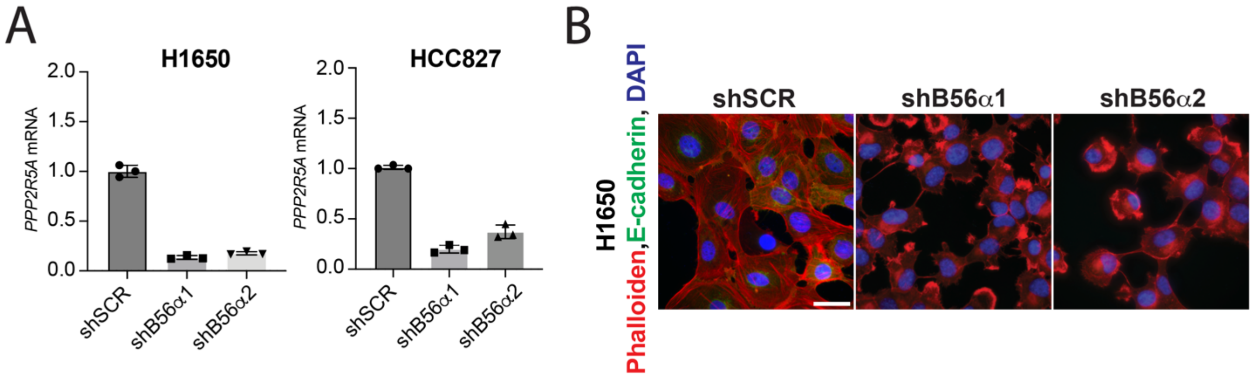


**Supplemental Figure 1:** A) mRNA expression of B56α (*PPP2R5A*) in H1650 and HCC827 shB56α cells compared to shSCR control. B) Merged representative images of H1650 cells with immunofluorescence for DAPI (blue), Phalloidin (F-actin, red), and E-cadherin (green)(scale bar = 100μm).
