## Supplemental Figure 2 for "Suppression of PP2A-B56α Drives EMT in EGFR Mutant Non-Small Cell Lung Cancer"

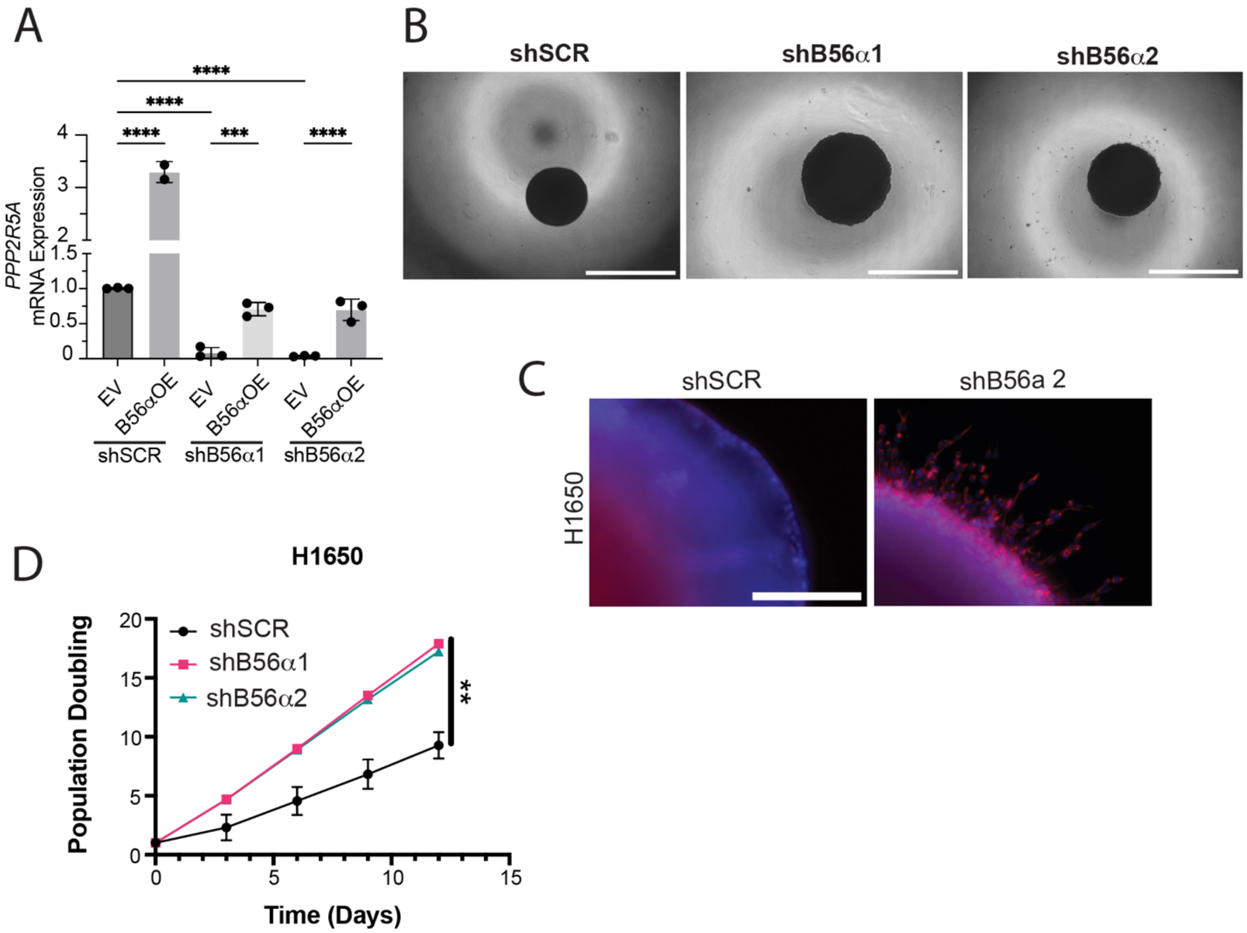


**Supplemental Figure 2:** A) mRNA expression of B56α (*PPP2R5A*) at the time of plating the transwell migration assay in Figure 3A. B) Representative images of spheroids at the time of embedding the spheroids in Matrigel for the spheroid invasion assay in Figure 3C (scale bar = 1mm). C) Representative immunofluorescent images of DAPI and Phalloidin (F-actin) in H1650 shB56α compared to shSCR (scale bar = 250μm). D) Population doubling of H1650 shB56α cells compared to shSCR control.
