## Supplemental Figure 3 for "Suppression of PP2A-B56α Drives EMT in EGFR Mutant Non-Small Cell Lung Cancer"

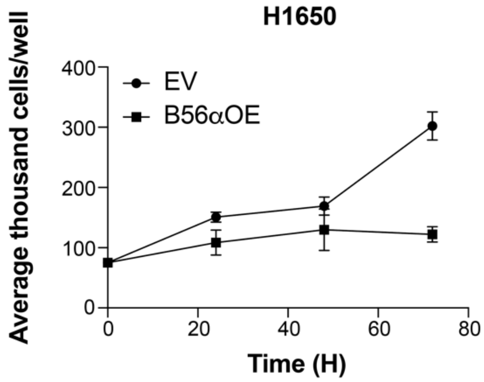


**Supplemental Figure 3:** Proliferation of stable overexpression of B56α (B56αOE) in H1650 cell line compared to empty vector control (EV). Shown as 3 technical replicates.
