## Supplemental Figure 4 for "Suppression of PP2A-B56α Drives EMT in EGFR Mutant Non-Small Cell Lung Cancer"

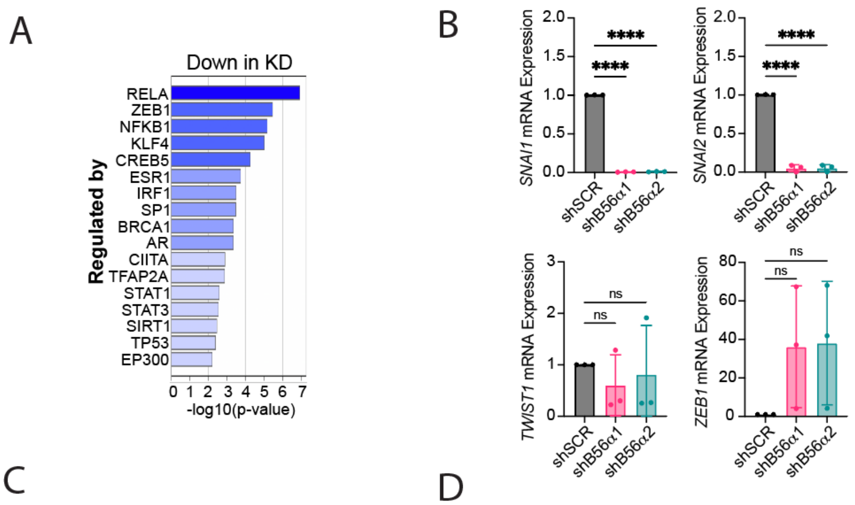


C

| Color | MCODE | GO | Description | Log10(P) |
|---|---|---|---|---|
| | MCODE_1 | GO:0002934 | desmosome organization | -18.9 |
| | MCODE_1 | R-HSA-6809371 | Formation of the cornified envelope | -15.3 |
| | MCODE_1 | R-HSA-6805567 | Keratinization | -13.6 |
| | MCODE_2 | WP383 | Striated muscle contraction pathway | -10.5 |
| | MCODE_2 | hsa04820 | Cytoskeleton in muscle cells | -9.8 |
| | MCODE_2 | GO:0007015 | actin filament organization | -9.4 |
| | MCODE_3 | M184 | PID ECADHERIN KERATINOCYTE PATHWAY | -12.1 |
| | MCODE_3 | GO:0044331 | cell-cell adhesion mediated by cadherin | -11.3 |
| | MCODE_3 | M156 | PID ECADHERIN NASCENT AJ PATHWAY | -10.9 |
| | MCODE_4 | GO:0034340 | response to type I interferon | -10.2 |
| | MCODE_4 | GO:0051607 | defense response to virus | -10.1 |
| | MCODE_4 | R-HSA-909733 | Interferon alpha/beta signaling | -10.0 |
| | MCODE_5 | R-HSA-432720 | Lysosome Vesicle Biogenesis | -12.1 |
| | MCODE_5 | R-HSA-199992 | trans-Golgi Network Vesicle Budding | -10.9 |
| | MCODE_5 | R-HSA-432722 | Golgi Associated Vesicle Biogenesis | -7.9 |
| | MCODE_6 | GO:0045109 | intermediate filament organization | -7.2 |
| | MCODE_6 | GO:0031424 | keratinization | -7.1 |
| | MCODE_6 | GO:0045104 | intermediate filament cytoskeleton organization | -6.9 |
| | MCODE_7 | GO:0045109 | intermediate filament organization | -7.8 |
| | MCODE_7 | GO:0045104 | intermediate filament cytoskeleton organization | -7.5 |
| | MCODE_7 | GO:0045103 | intermediate filament-based process | -7.5 |

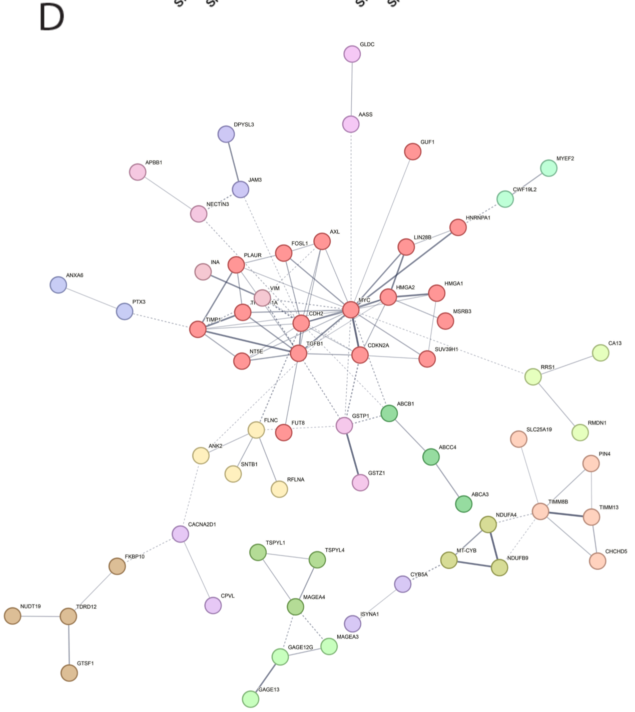


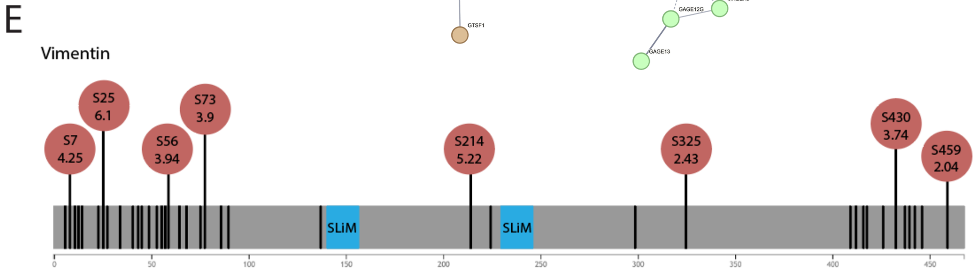


**Supplemental Figure 4:** A) TRRUST analysis of transcription factor target interactions that are decreased in shB56α total proteomics compared to shSCR control ($Log_2FC < -2$, $p<0.05$). B) mRNA expression analysis by qRT-PCR of classical EMT-TFs including SNAIL (*SNAI1*), SLUG (*SNAI2*), TWIST (*TWIST1*), and ZEB1 (*ZEB1*) in shB56α compared to shSCR cells (n=3 biological replicates). C) MCODE clusters from the Metascape MCODE analysis of significantly decreased proteins in the total proteomics ($Log_2FC < -2$, $p<0.05$). D) STRING enrichment analysis for top 15% of differentially upregulated proteins in shB56α cells. E) Vimentin protein phosphorylation sites denoted by black lines with changed phosphorylation indicated in red circles with the $Log_2FC$. Predicted B56 binding domains (SLiMs) shown in blue.
