## Supplemental Figure 5 for "Suppression of PP2A-B56α Drives EMT in EGFR Mutant Non-Small Cell Lung Cancer"

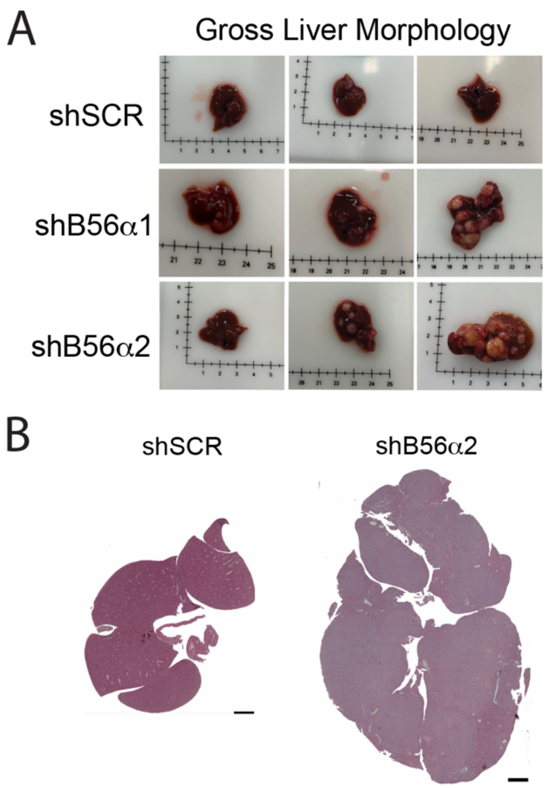


**Supplemental Figure 5:** A) Gross morphology of liver from mice injected with H1650 shB56α or shSCR control cells. B) H&E of liver showing increased tumor in shB56α2 compared to shSCR (scale bar = 2mm).
