## Supplemental Methods Table 1 for "Suppression of PP2A-B56α Drives EMT in EGFR Mutant Non-Small Cell Lung Cancer"

**Supplemental Methods Table 1: qRT-PCR primer sequences for genes analyzed.**

| Gene Name | Species | Forward Sequence | Reverse Sequence |
|---|---|---|---|
| ***PPP2R5A*** **B56α** | Human | 5'-AgAgCCCTgATTTCCAgCCTA-3'<br>5'-CTTTgCATTgCCACTgAAAg-3' | 3'-TTTCCCATAAATTCggTgCAgA-5'<br>3'-CAgCAgTCCTCTgATCAC-5' |
| ***CDH1*** | Human | 5'-gAACgCATTgCCACATAC-3' | 3'-ACCTTCCATgACAgACCC-5' |
| ***VIM*** | Human | 5'-AgTCCACTgAgTACCggAgAC-3' | 3'-CATTTCACgCATCTggCgTTC-5' |
| ***MYC*** | Human | 5'-CAAACCTCCTCACAgCCCACT-3' | 3'-TTCgCCTCTTgACATTCTCCTC-5' |
| ***KIAA1524*** **(CIP2A)** | Human | 5'-gCCACACTgATTCggTgTTTT-3' | 3'-TgCCgACAAAgATTTgCCAATA-5' |
| ***ZEB1*** | Human | TTACACCTTTgCATACAgAACCC | TTTACgATTACACCCAgACTgC |
| ***TWIST*** | Human | 5'-CCAggTACATCgACTTCCTCTA-3' | 3'-CCATCCTCCAgACCgAgAA-5' |
| ***SNAIL*** | Human | ACTgCAACAAggAATACCTCAg | gCACTggTACTTCTTgACATCTg |
| ***SLUG*** | Human | TgTgACAAggAATATgTgAgCC | TgAgCCCTCAgATTTgACCTg |
| **18s** | Human | 5'-CACCAACATCgATgggCgg-3' | 3'-CACACgTTCCACCTCATCCTCAg-5' |
