## Supplemental Methods Table 2 for "Suppression of PP2A-B56α Drives EMT in EGFR Mutant Non-Small Cell Lung Cancer"

**Supplementary Methods Table 2: Antibodies used for western blot analysis.**

| Antibody | Catalog Number | LOT Number | Company | Dilution |
|---|---|---|---|---|
| **GAPDH** | 4300 | 01062548 | Fisher | 1:20,000 |
| **pEGFR (Y1068)** | 2236 | 18 | Cell Signaling Technology | 1:1000 |
| **EGFR** | 2239 | 5 | Cell Signaling Technology | 1:1000 |
| **Cleaved PARP** | 5625 | 18 | Cell Signaling Technology | 1:1000 |
| **c-MYC** | ab32072 | 1053331-27 | Abcam | 1:1000 |
| **CIP2A** | 80659 | H2522 | Santa Cruz Biotechnology | 1:1000 |
| **E-cadherin** | 14472S | 8 | Cell Signaling Technology | 1:1000 |
| **pS6 (S240/244)** | 5364 | 13 | Cell Signaling Technology | 1:1000 |
| **S6** | 2317 | 13 | Cell Signaling Technology | 1:1000 |
| **Goat anti-Mouse IgG Secondary Antibody** | NC9401841 | D20802-25 | LICOR | 1:5000 |
| **Goat anti-Rabbit IgG Secondary Antibody** | NC0252291 | D20809-05 | LICOR | 1:5000 |
