## Supplemental Methods Table 3 for "Suppression of PP2A-B56α Drives EMT in EGFR Mutant Non-Small Cell Lung Cancer"

**Supplemental Data Methods 3: Antibodies used for immunofluorescence *in vitro* and tissue analysis.**

| Antibody | Catalog Number | LOT Number | Company | Dilution |
|---|---|---|---|---|
| Mouse E-cadherin | 610181 | 3138351 | BD Biosciences | 1:200 |
| Rabbit Vimentin | 5741S | 8 | Cell Signaling Technology | 1:200 |
| Rhodamine Phalloidin | R415 | 2641902 | Invitrogen | 1:200 (*in vitro*) |
| Ku-80 | 2180S | 3 | Cell Signaling Technology | 1:600 (tissue) |
| Goat anti-Rabbit IgG (H+L) Alexa Fluor™ Plus 594 | A32740 | VA295503 | Invitrogen | 1:2000 (*in vitro*)<br>1:1000 (tissue) |
| Goat anti-Mouse IgG (H+L) Alexa Fluor™ Plus 594 | A32742 | VA295504 | Invitrogen | 1:2000 (*in vitro*)<br>1:1000 (tissue) |
| Goat anti-Rabbit IgG (H+L) Alexa Fluor™ Plus 488 | A32731 | VA295501 | Invitrogen | 1:2000 (*in vitro*)<br>1:1000 (tissue) |
| Goat anti-Mouse IgG (H+L) Alexa Fluor™ Plus 488 | A32723 | VA297822 | Invitrogen | 1:2000 (*in vitro*)<br>1:1000 (tissue) |
| Goat anti-Rabbit IgG (H+L) Alexa Fluor™ Plus 647 | A32733 | VB296618 | Invitrogen | 1:2000 (*in vitro*)<br>1:1000 (tissue) |
| Goat anti-Mouse IgG (H+L) Alexa Fluor™ Plus 647 | A32728 | UK290265 | Invitrogen | 1:2000 (*in vitro*)<br>1:1000 (tissue) |
